# Identifying Sialylation Linkages at the Glycopeptide Level by Glycosyltransferase Labeling Assisted Mass Spectrometry (GLAMS)

**DOI:** 10.1101/811554

**Authors:** He Zhu, Shuaishuai Wang, Ding Liu, Lang Ding, Congcong Chen, Yunpeng Liu, Zhigang Wu, Roni Bollag, Kebin Liu, Jun Yin, Cheng Ma, Lei Li, Peng George Wang

## Abstract

Precise assignment of sialylation linkages at the glycopeptide level is of importance in bottom-up glycoproteomics, and is also an indispensable step to understand the function of glycoproteins in pathogen-host interactions and cancer progression. Even though some efforts have been dedicated to the discrimination of α2,3/α2,6-sialylated isomers, unambiguous identification of sialoglycopeptide isomers is still needed. Herein, an innovative strategy of glycosyltransferase labeling assisted mass spectrometry (GLAMS) was developed. After specific enzymatic labeling, oxonium ions from higher-energy C-trap dissociation (HCD) fragmentation of α2,3-sailoglycopeptides generate unique reporters to distinctly differentiate those of α2,6-sailoglycopeptide isomers. Using this strategy, a total of 1,236 linkage-specific sialoglycopeptides were successfully identified from 161 glycoproteins in human serum.

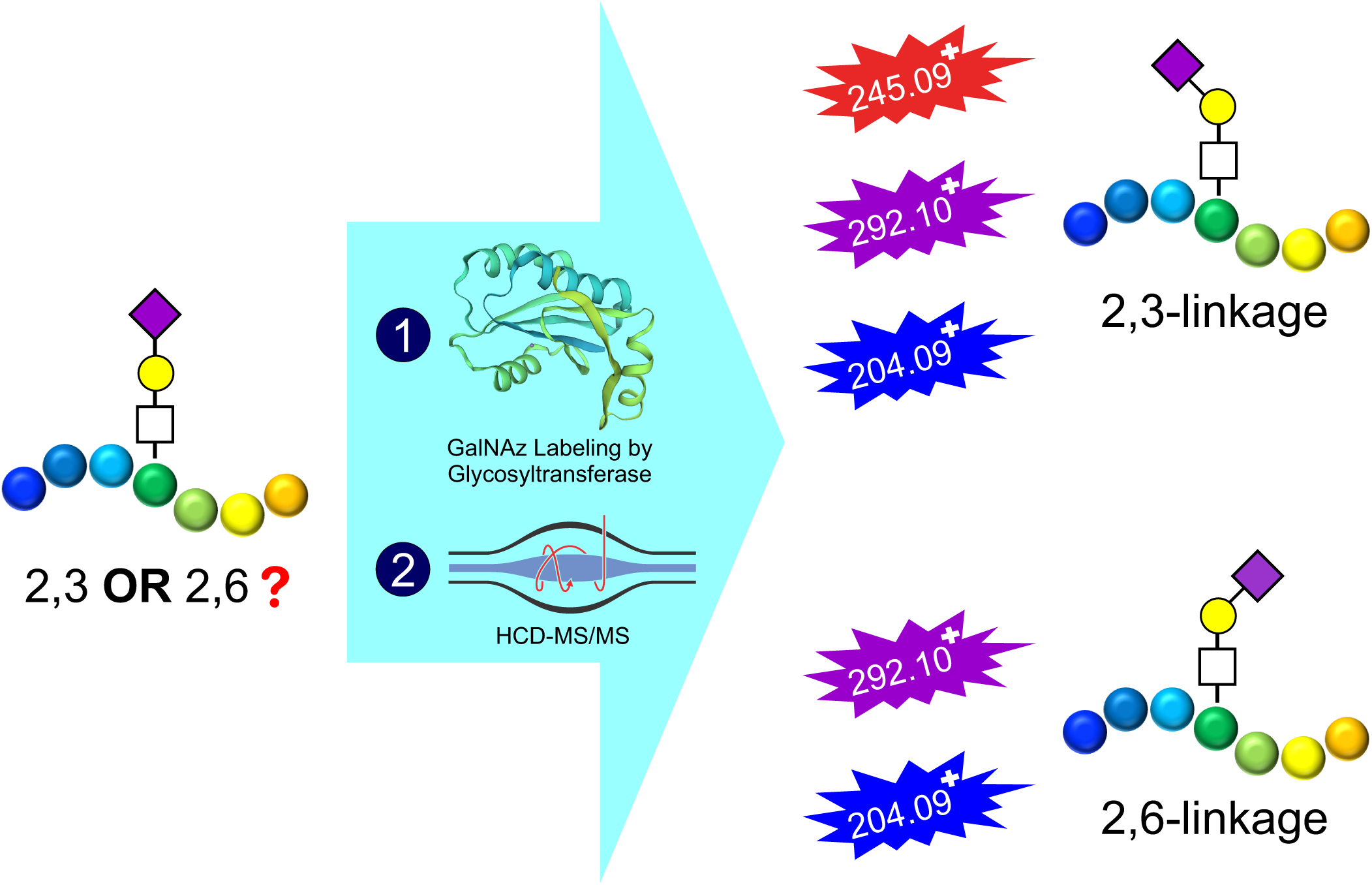

## INTRODUCTION

Glycosylation is one of the most important post-translational modifications (PTMs) of proteins^1^. It plays an indispensable role in a wide variety of biological and physiological processes including protein folding^2^, cellular communication and signaling^3^, immune response^4^, and pathogen-host interactions^5^. It was estimated that more than half of proteins in the human proteome are glycosylated^6^. Glycosylation is generally classified into N-glycosylation and mucin-type O-glycosylation. In N-glycosylation, the glycans are attached to asparagine residues of proteins in the consensus motif Asn-X-Ser/Thr/Cys (X can be any amino acid except for proline)^7^, whereas the glycan moieties are linked to serine or threonine residues of proteins in O-glycosylation^1^. N-acetylneuraminic acid (Neu5Ac) is most common form of diverse sialic acids found on glycan chains in humans^8^. Neu5Ac is the frequently the terminal monosaccharide residue that linked to galactose (Gal) via an α2,3 or an α2,6 linkage^9^. The two Neu5Ac linkage isomers have important roles in pathogen binding. For example, avian influenza A viruses preferentially bind to respiratory epithelial cells that contain α2,3-linked Neu5Ac moieties, while human influenza viruses selectively bind to α2,6-linked Neu5Ac moieties^10^. In addition, altered sialylation linkages is closely correlated with health status. In patients diagnosed with prostate cancer, the structure of sialylated glycans released from the prostate-specific antigen (PSA) is rich in α2,3 linkages, whereas the structure of sialylated glycans from healthy people’s PSA is predominantly via α2,6 linkages^11^. Without detailed information on sialylation linkages, it is challenging to decipher the biological functions of sialoglycoproteins.

A variety of methods, including sialidase treatment^12^, chemical derivatization^13–21^, and mass spectrometry (MS)^22–24^, have been used to distinguish α2,3 and α2,6-sialylated glycans, but the location of corresponding glycosylation sites is not evident. The emerging technology of MS-based glycoproteomics^25–29^ provides a high-throughput approach to analyze site-specific protein glycosylation. Consequently, strategies for identifying sialylation linkages at the glycopeptide level are required. Established chemical approaches^30, 31^ modify not only sialic acids of the glycan moiety but also acidic amino acid residues and the C-terminus of the peptide backbone, which increases the complexity of spectra analysis, especially in relation to large scale glycoproteomics data^32^. Pseudo MS^3^ fragmentation^22^ and ion mobility-mass spectrometry (IM-MS)^33^ have also been applied to distinguish sialoglycopeptide isomers but this technology has not been **demonstrated (**?effective) for large-scale glycopeptides analysis. Even though the fragmentation of high energy C-trap dissociation (HCD) can achieve the high-throughput assignment of α2,3/α2,6-sialoglycopeptide isomers, there are still many ambiguous glycopeptides without sialylation linkage information^34^.

Enzymatic labeling has become a promising tool for probing proteins with specific glycosylation because of its high specificity and mild reaction condition^35–41^. In our previous research, a β1,4-N-acetylgalactosaminyltransferase from *Campylobacter jejuni* (CgtA) was demonstrated to specifically recognize the Neu5Ac(α2,3)Gal epitope and transfer N-acetylgalactosamine (GalNAc) or N-azidoacetylgalactosamine (GalNAz) to Gal residue via β1,4 linkage^42^. Taking advantage of the specificity of CgtA, we herein report a new strategy for identification of α2,3 and α2,6 sialoglycopeptide isomers by glycosyltransferase labeling assisted mass spectrometry (GLAMS). As shown in Scheme 1, sialoglycopeptides (monosialylation for example) without linkages information were treated with GalNAz labeling by CgtA. After HCD fragmentation, if only the oxonium ions of HexNAc (N-acetylhexosamine, m/z 204) and Neu5Ac (m/z 292) are observed in the spectrum, this identifies an α2,6 linkage; if one more oxonium ion m/z 245 from GalNAz is included, then it represents an α2,3 linkage. We demonstrated the capacity of GLAMS to effectively differentiate α2,3/α2,6 isomers using synthetic sialoglycopeptides, and applied it to profile sialylated N-glycopeptides digested from serum samples, in which we identified 1,236 linkage specific sialoglycopeptides with an improved version of pGlyco 2.0 software^27^.

**Scheme 1.**
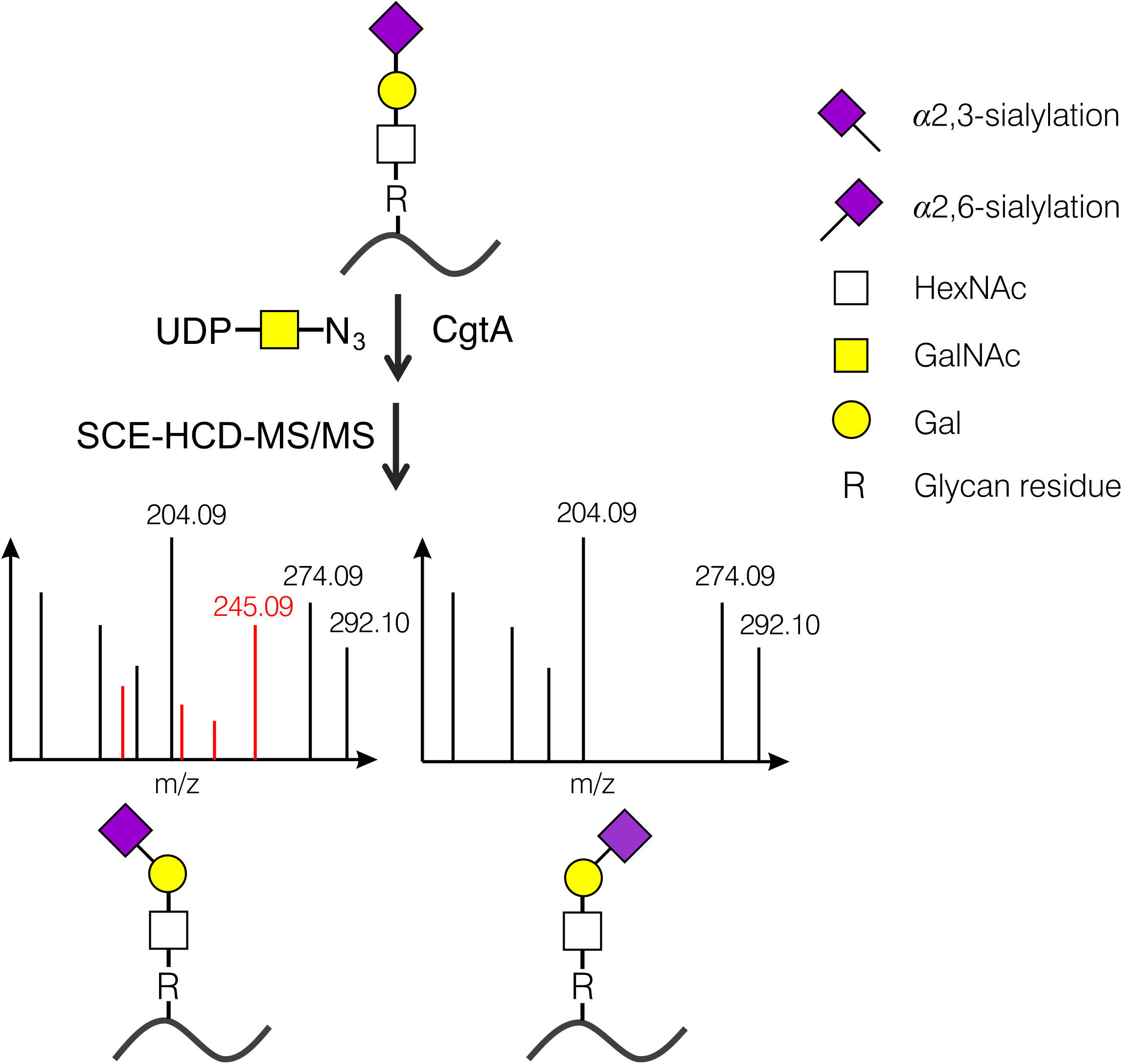
Strategy for distinguishing α2,3/α2,6-sialoglycopeptides by glycosyltransferase labeling assisted mass spectrometry.

## EXPERIMENTAL SECTION

### Chemicals and materials

Amicon Ultra-0.5 Centrifugal Filter Unit with NMWL of 30 kD was obtained from Millipore Sigma (St. Louis, MO). iSPE^®^ HILIC SPE Cartridge was purchased from The Nest Group (Southborough, MA). XAmide medium was provided by Dr. Xinmiao Liang’s lab (Dalian Institute of Chemical Physics, Chinese Academy of Sciences). Sequencing grade porcine trypsin was purchased from Promega (Madison, WI); Deionized water was produced by a Milli-Q A10 system from Millipore (Bedford, MA). Optimal LC-MS grade quality acetonitrile (ACN), formic acid (FA) and water were purchased from Thermo Fisher Scientific, Inc. (Waltham, MA). Dithiothreitol (DTT), 2-Iodoacetamide (IAA) and trifluoroacetic acid (TFA) were bought from Alfa Aesar (Ward Hill, MA). Ammonium bicarbonate (NH_4_HCO_3_) was purchased from Millipore Sigma (St. Louis, MO). All other chemicals of analytical grade were commercially available.

Sialylated oligosaccharide standards including 3’-sialyllactose (3SL) and 6’-sialyllactose (6SL) were enzymatically synthesized using *Pasteurella multocida* α2,3-sialyltransferase 1 and *Photobacterium damselae* α2,6-sialyltransferase as described before^43^.

Serum samples from four healthy people, ten patients with colon cancer and four patients with liver cancer were provided by Georgia Cancer Center at Augusta University and stored at - 80 °C until use. The protocol for serum sample preparation was approved by the Institutional Review Board of Augusta University and was performed in accordance with the Helsinki Declaration. All participants gave written informed consents.

### Protein digestion

The protein digestion was performed using a filter aided sample preparation (FASP)^44^ method with minor modification. Approximately 1 mg of total proteins from human serum was subjected to FASP procedures. Samples were denatured in 20 μL of lysis buffer (4% SDS, 100 mM DTT, 20 mM Tris-HCl, pH 7.6) at 95 °C for 10 min. When the solution was cooled down to room temperature, 1 mL of UA solution (8 M urea, 20 mM Tris-HCl, pH 8.5) was added. The solution was gradually transferred into a 30 kD filter and centrifugated at 15,000 rpm at 20 °C for 15 min. Then 100 μL of IAA solution (50 mM IAA in UA solution) was added to the filter and incubated for 30 min in the darkness followed by centrifugation at 15,000 rpm for 10 min. After that, 100 μL of UA solution and 200 μL of digestion buffer (100 μL of 50 mM NH_4_HCO_3_ each time) were added and centrifuged at 15,000 rpm for 10 min, respectively. Finally, 20 μg of trypsin was added into the filter and incubated at 37 °C for 12 h. The tryptic peptides were eluted using 100 μL of 50 mM NH_4_HCO_3_ for six times. The concentration of the collected peptides was measured based on absorbance at 280 nm with Nanodrop. Five hundred µg of the peptides were taken and subsequently dried in Vacufuge at room temperature, then were lyophilized twice to remove NH_4_HCO_3_. The dried samples were stored at −20 °C for further use.

### Enrichment of glycopeptides

The enrichment of glycopeptides was performed using an iSPE HILIC cartridge by the procedure of ZIC-HILIC^45^. Briefly, the HILIC cartridge was pre-washed with 300 µL of 0.1% TFA and conditioning with 600 µL of 80% ACN containing 0.1% TFA. Peptide samples dissolved in 400 µL of 80% ACN containing 0.1% TFA were loaded onto the cartridge. The flow-through was reloaded onto the column for another two times. Then the column was washed with 1.2 mL of 80% ACN containing 1% TFA. The glycopeptides were eluted using 750 µL of 0.1% TFA, 60 µL of H_2_O, 60 µL of 25 mM NH_4_HCO_3_ and 60 µL of 50% ACN. The fractions were combined followed by lyophilization and stored at −20 °C.

### Enzymatic labeling of glycopeptides

The β1,4-N-acetylgalactosaminyltransferase from *C. jejuni* (CgtA) and sugar nucleotide UDP-N-azidoacetylgalactosamine (UDP-GalNAz) were prepared as described previously^42^. For enzymatic labeling, the 40 µL reaction mixture containing the enriched glycopeptides, 50 mM Tris-HCl buffer (pH 7.5), 5 mM MgCl_2_, 200 µM UDP-GalNAz and 10 µg CgtA was incubated in a 37 °C water bath for 1 h. The reaction was quenched by adding 60 µL of ACN, 6 µL of 10% TFA and 215 µL of ACN containing 1% TFA. Then the glycopeptides were purified with an in-house packed Amide HILIC microcolumn (6 mg of XAmide HILIC medium in a 200 µL tip) which was preconditioned with 100 µL of 0.1% TFA and 200 µL of 85% ACN containing 0.1% TFA. After loading the reaction mixture onto the tip column, the column was washed with 100 µL of 85% ACN, 0.1% TFA to remove salts and proteins. The glycopeptides were collected using 200 µL of 0.1% TFA and lyophilized.

### Nano LC-MS/MS analysis

Nano RP HPLC-MS experiments were performed on an LTQ-Orbitrap Elite mass spectrometer (Thermo Fisher) equipped with EASY-spray source and nano-LC UltiMate 3000 high-performance liquid chromatography system (Thermo Fisher). The separation was performed on an EASY-Spray PepMap C18 Column (75 μm × 50 cm, 2 μm, ThermoFisher, US) with a linear gradient from 3% to 40% mobile phase B for 120 min at a flow rate of 300 nL/min (mobile phase A: 1.95% ACN, 97.95% H_2_O, 0.1% FA; mobile phase B: 79.95% ACN, 19.95% H_2_O, 0.1% FA). The mass spectrometer was operated in the data-dependent mode. A full-scan survey MS experiment (m/z range from 375 to 2000; automatic gain control target, 1,000,000 ions; resolution at 400 m/z, 60,000; maximum ion accumulation time, 50 ms) was acquired by the Orbitrap mass spectrometer, and ten most intense ions were fragmented by higher-energy C-trap dissociation (HCD) with an optimized stepped collision energy (SCE) of 30 ± 15% NCE.

### Data analysis

The pGlyco 3.0 software (an improved version of pGlyco 2.0^27^) was used to identify N-glycopeptides. In the glycan database, we defined an unnatural monosaccharide GalNAz (Gaz, Z). We created all the possible “Gaz” containing human N-glycan sequences following the motif H(A)(Z), where “H” represents Galactose, “A” represents Neu5Ac and “Z” represents GalNAz. For the sialoglycopeptides identification, the raw data was first converted to an mgf file using pParse^46^. The potential sialoglycopeptides spectra containing oxonium ions of both GlcNAc (m/z 204.09, m/z 186.08) and Neu5Ac (m/z 292.10, m/z 274.09) were extracted and merged into a new mgf file using an in-house program. The new mgf file was searched against an Uniprot-Swiss human protein database (20,417 reviewed entries on March 20, 2019) and a modified N-glycan database. Other searching parameters were used including carbamidomethyl (Cys) as a fixed modification, deamination (Asn) and oxidation (Met) as variable modifications, and trypsin as the digestion enzyme with maximum two missed cleavage sites allowed. The mass tolerance for the precursor ions and the fragment ions was set to 10 ppm and 20 ppm, respectively. A total false discovery rate (FDR) of 1% was applied to all data sets at the glycopeptide level. The identified glycoproteins were annotated using Uniprot ID mapping^47^.

## RESULTS AND DISCUSSION

### The specificity of CgtA for Sialoglycopeptides

Our previous study showed that CgtA had a high specificity for (Neu5Ac**-**α2,3-Gal)-containing glycans^42^. To test its selectivity for sialoglycopeptides, we synthesized 23SP and 26SP (synthesis and data not published) as shown in Figure 1. The activity of CgtA was validated with sialolactose prior to the enzymatic reaction of sialoglycopeptides. After labeling with GalNAz by CgtA, only α2,3-sialylated lactose showed a mass shift of 244 Da in the MS spectrum and no substrate was obviously observed (**Figure S1**), which indicated that CgtA had good catalytic activity and high specificity as well.

**Figure 1.**
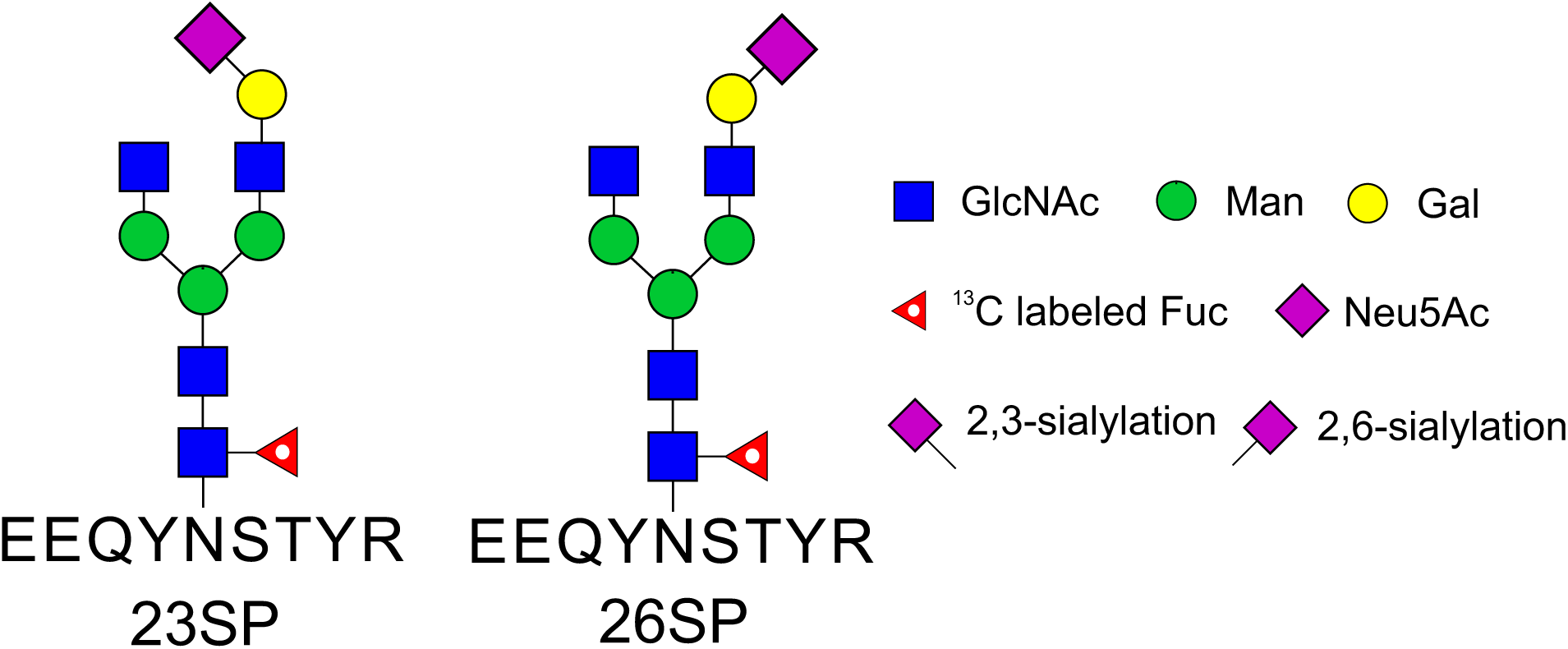
Structure of the synthesized α2,3-sialoglycopeptides (23SP) and α2,6-sialoglycopeptides (26SP).

ESI-MS was used to monitor the GalNAz labeling reaction for sialoglycopeptides. For the α2,3-sialoglycopeptide 23SP, the monoisotopic peak at m/z 1032.0808 is from the triply charged glycopeptide (**Figure S2A**). After incubating with CgtA and UDP-GalNAz for 30 min, a new dominant peak at m/z 1113.4422 with three positive charges was observed, which correspond to the GalNAz labeled 23SP (**Figure S2C**). However, we could still detect minor substrate 23SP. The α2,6-sialoglycopeptide 26SP has the same exact mass (**Figure S2B**) as 23SP. After GalNAz labeling by CgtA, the strongest peak is located at the same exact position, and a trace amount of product at m/z 1113.4422 was detected (**Figure S2D**). This may result from the very weak activity of CgtA for 2,6-sialylated glycans^42^.

The reaction time was further optimized to increase the conversion rate of GalNAz labeling of 23SP. The reaction mixture at different time points was analyzed by LC-MS and the reaction efficiency was calculated based on the peak area. For 23SP, in the first half an hour, the reaction efficiency was dramatically increased. After that, the increase slowed down and the reaction efficiency approached 100% at 60 min. Meanwhile, the conversion rate for the 26SP remained unchanged (less than 1%) over time (Figure 2). Finally, we chose 60 min as the reaction time for the subsequent GalNAz labeling reactions.

**Figure 2.**
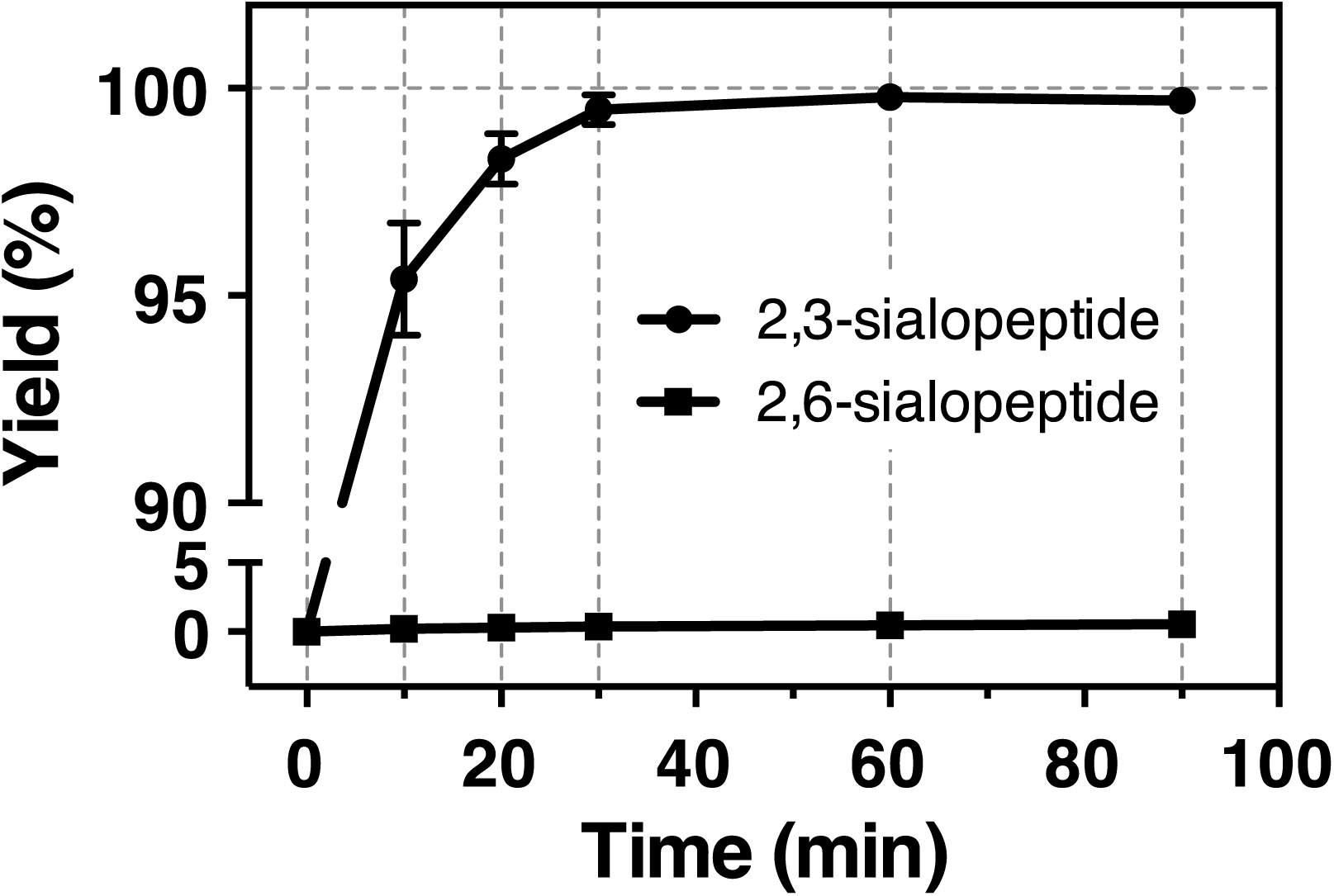
Time optimization for CgtA labeling of 23SP and 26SP.

### Identifying Sialoglycopeptide Linkage Isomers by GalNAz Labeling and Tandem MS

Although discrimination of α2,3/α2,6-sialoglycopeptide isomers can be achieved by the mass shift in primary MS as shown above, we cannot accurately determine the glycopeptide sequence using only molecular weight information. Tandem MS is an effective way for sequencing glycopeptides, and HCD-MS/MS is commonly used to analyze glycopeptides because of the high accuracy and detectable diagnostic oxonium ions of monosaccharides^26–28, 48^. It was also demonstrated that HCD with stepped collision energy (SCE) can simultaneously determine the sequences of both glycans and peptides in a single run^27, 48^. To achieve a better sequence coverage of glycopeptides on the Orbitrap we used, SCE was optimized with glycopeptides enriched from human serum. The identification of N-glycopeptides was performed using pGlyco 2.0 software^27^. A series of two-step SCEs and three-step SCEs were employed to compare the number of identified glycopeptides. As shown in **Figure S3**, with a three-step SCE of “15-30-45”, more glycopeptides can be identified from the same sample.

The optimized SCE-HCD-MS/MS was applied for sequencing 23SP and 26SP. After GalNAz labeling, 23SP (Figure 3A) showed diagnostic oxonium ions of GlcNAc and Neu5Ac at m/z 204.09 and 292.10, which are common features of sialoglycopeptides. Apart from these oxonium ions, 23SP also had a strong unknown peak at m/z 245.09. Compared with the spectrum of 23SP before GalNAz labeling (**Figure S4**), we can obviously see that the peak with m/z 245.09 corresponds to the oxonium ion of GalNAz. Following the dissociation pathway of GlcNAc^39, 49, 50^ in Scheme S1A, a series of fragmentation ions of GalNAz in HCD were illustrated in Scheme S1B. The peaks at m/z 245.09, m/z 227.08, m/z 209.07 and m/z 179.06, which match the theoretical values very well, are diagnostic ions of GalNAz (**Figure S5**). For 26SP (Figure 3B), its spectrum showed that it shared similar fragmentation patterns with GalNAz labeled 23SP. In the low m/z region, only diagnostic ions of GlcNAc and Neu5Ac were observed, which clearly demonstrated that the GLAMS is an effective approach to distinguish intact linkage specific sialoglycopeptides.

**Figure 3.**
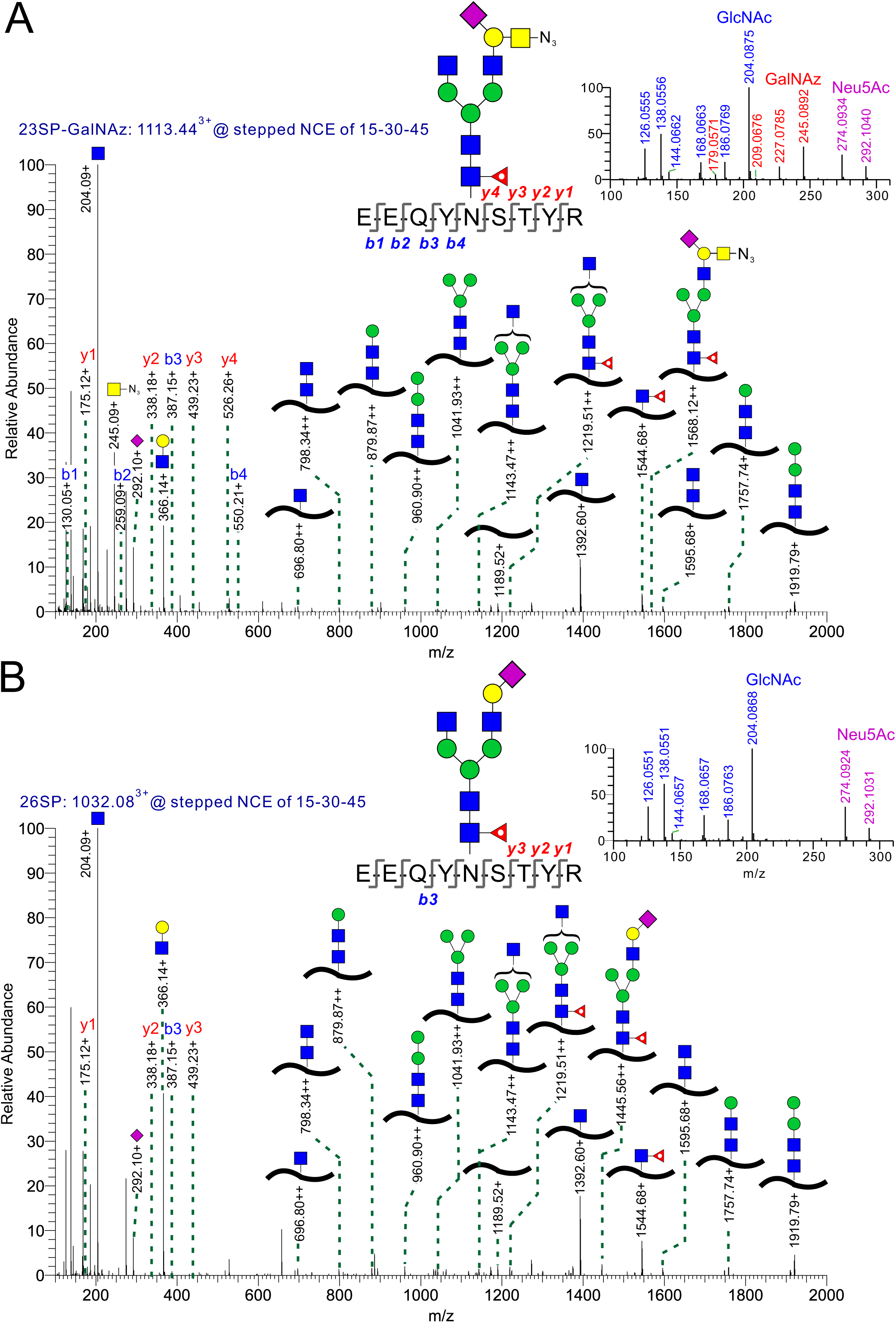
HCD-MS/MS spectra of 23SP (A) and 26SP (B) after enzymatic labeling.

### Linkage Specific Sialoglycopeptides in Human Serum

We then applied this method to analyze the N-linked α2,3/α2,6-sialoglycopeptides of complex biological samples. Serum samples from healthy individuals (n = 4) and patients diagnosed with liver cancer (n = 4) and colon cancer (n = 10) were used in this study. In the beginning, we tried to perform the enzymatic labeling of the mixture of tryptic peptides and glycopeptides. After drying from the FASP eluents, the mixture was not completely dissolved in the Tris buffer (pH 7.5) for the enzymatic reaction, which may result from the existence of some basic peptides. To solve the solubility issue, we adopted step-by-step glycopeptides enrichment and enzymatic labeling as shown in the workflow Figure 4. It turned out that the enriched glycopeptides were dissolved in the basic buffer very well. This may be attributed to that conjugated glycans increase the hydrophilicity of peptides and some sialoglycans decrease the isoelectric points of peptides according to the previous findings^51, 52^.

**Figure 4.**
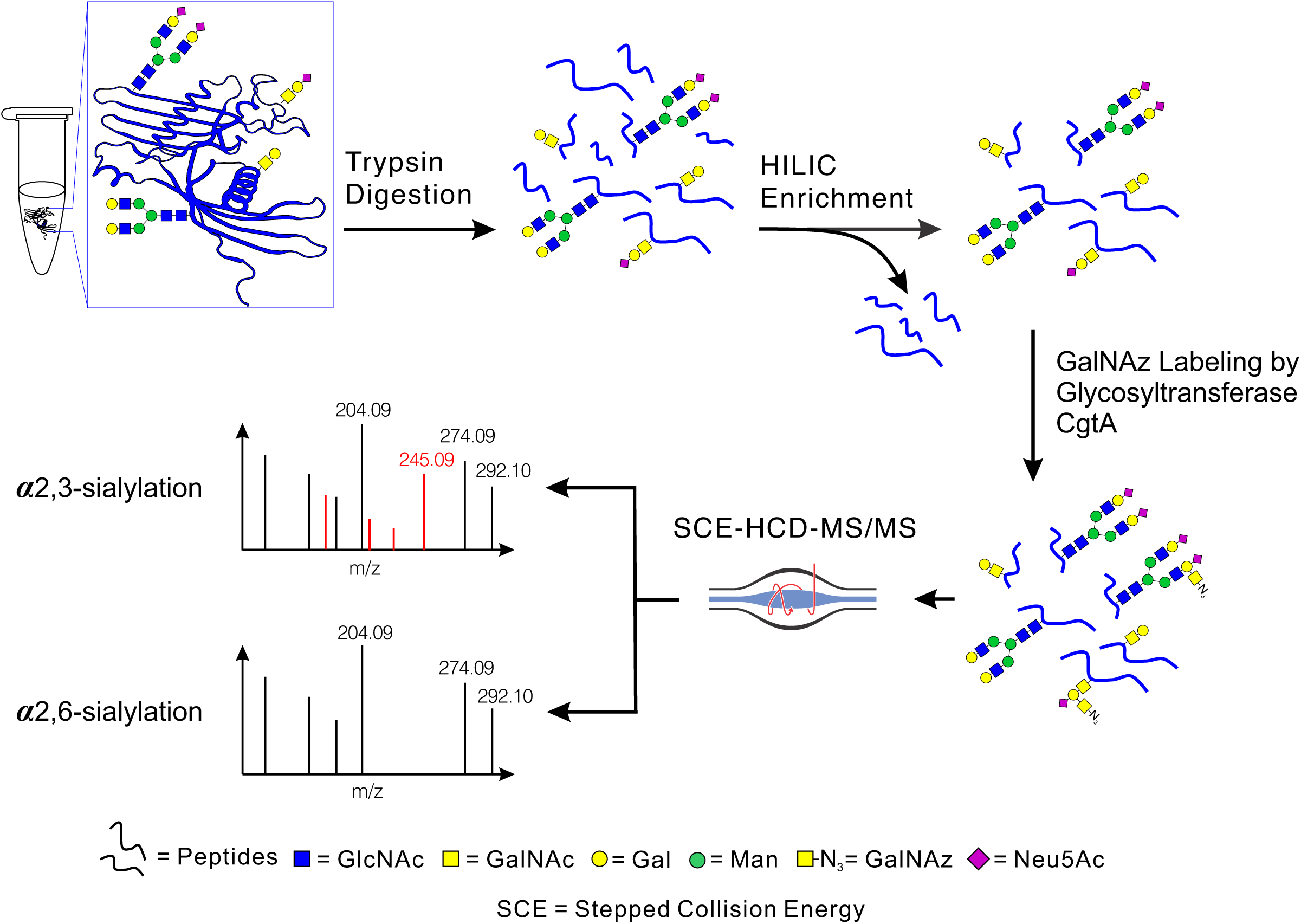
Workflow for identification of linkage specific sialoglycopeptides from human serum.

All the serum samples were analyzed in three replicated LC-MS/MS runs for a large-scale study of N-linked sialoglycopeptides. Using the oxonium ions of GalNAz as a tag to mark α2,3-sialylation, a total of 22,819 sialoglycopeptide spectra were identified with a 1% total FDR, corresponding to 1,236 unique site-specific sialoglycans on 261 N-glycosylation sites from 161 unique glycoproteins (**Table S1**), among of which 53% was also identified previously using a deglycosylation method^53^ as shown in **Figure S6**. For the distribution analysis of α2,3 or α2,6-sialylation (Figure 5A), only 48 distinct site-specific sialoglycopeptides contained pure α2,3-sialylation linkage, 236 sialoglycopeptides contained hybrid sialylation, and all the rest were uniquely 2,6-sialylated, which accounts for more than 77% of the identified sialoglycopeptides. This is consistent with the previous result that α2,6-sialylation is dominant in sialylated N-glycans released from human serum samples^54^. Taking a closer look at the distribution of sialylation levels, monosialylation is the major component in strictly α2,3 or α2,6-sialoglycopeptides, and 74% of the hybrid sialoglycopeptides are disialylated.

**Figure 5.**
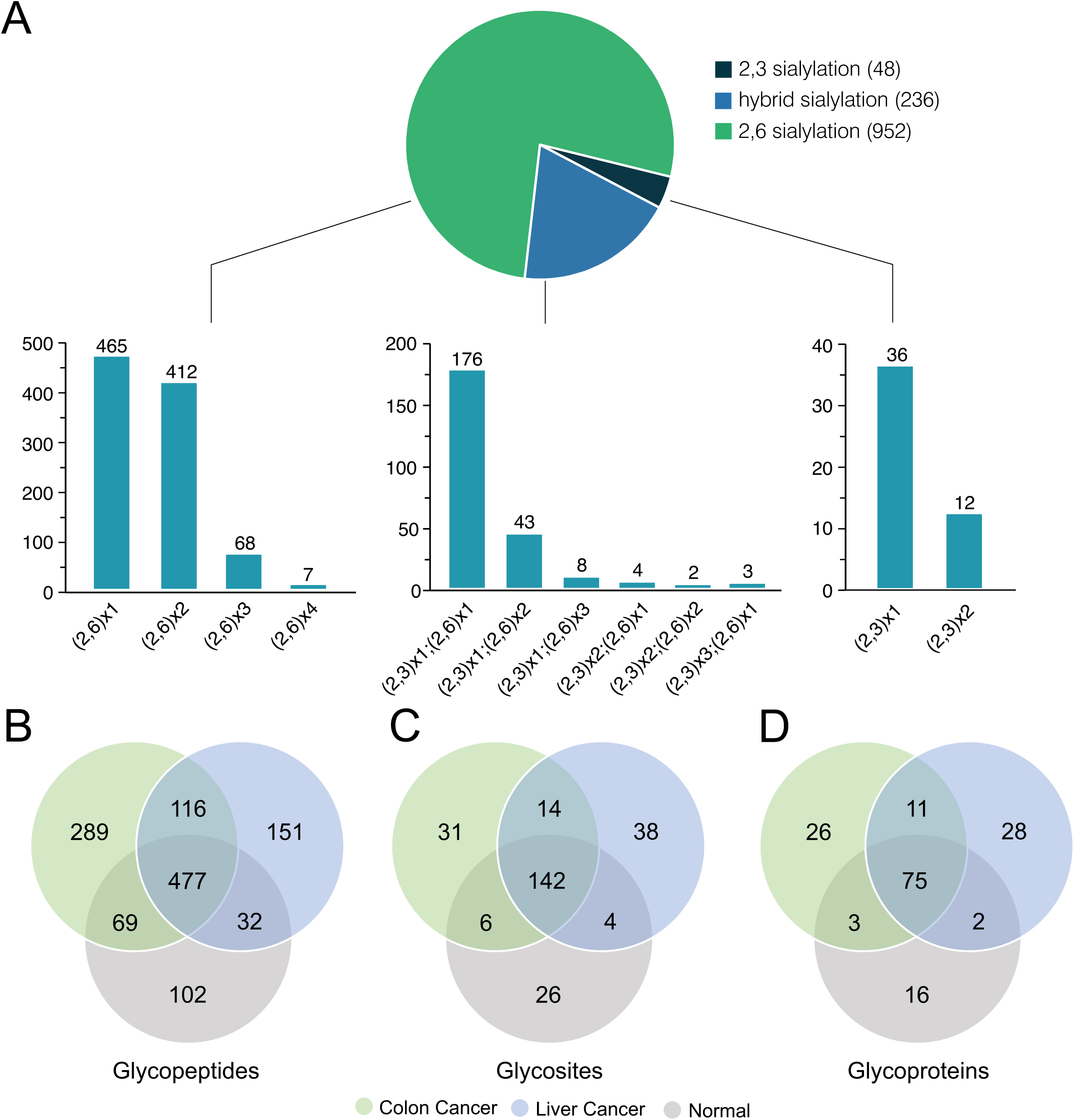
(A) Distribution of the sialoglycopeptides from human serum; Venn diagrams of identified glycopeptides (B), N-glycosylation sites (C), and glycoproteins (D) in serum samples from colon cancer patients, liver cancer patients and normal people.

The percentage of the overall overlapped glycosites (Figure 5C) and glycoproteins (Figure 5D) in a specific type of serum is 72-80% and 65-78%, respectively. The percentage of overlapping sialoglycopeptides (Figure 5B) in each kind of serum sample was significantly lower than those of the glycosites and glycoproteins, reflecting the diversity of site-specific glycosylation, especially for samples from colon cancer and liver cancer patients. A highly glycosylated protein P01876 (IGHA1) was selected to demonstrate the diversity of site-specific glycosylation in the three different sera. The previous study showed that IgA was α2,3 and α2,6-sialylated on N-glycans^55^. Here, we identified 35 unique sialoglycopeptides including α2,3 and α2,6 linkages on one N-glycosylation site of IGHA1 (**Table S2**). In addition, we found some unique sialoglycopeptides of IGHA1 expressed in a serum-specific manner. For example, only unique α2,6-sialoglycopeptides were identified in serum from liver cancer patients (representative spectrum in **Figure S7**) and normal people (representative spectrum in **Figure S8**), whereas unique α2,6-sialoglycopeptides and one hybrid sialoglycopeptide were detected from colon cancer patients’ sera (representative spectra in **Figure S9**).

## CONCLUSIONS

Elucidating linkage-specific sialylation of glycoproteins is significant to understand pathogen-host interactions and disease development. By integrating enzymatic labeling and MS-based glycoproteomics, we developed a high-throughput glycosyltransferase labeling assisted mass spectrometry (GLAMS) to identify sialoglycopeptide linkage isomers. Facilitated by the acceptor specificity of the glycosyltransferase CgtA toward the Neu5Ac(α2,3)Gal epitope, we have demonstrated the distinct diagnostic oxonium ions as tags to effectively discriminate α2,3- and α2,6-sialoglycopeptide isomers using LC separation coupled with SCE-HCD-MS/MS fragmentation. We also successfully applied this approach to a large-scale profiling of linkage-specific sialoglycopeptides in human serum samples. The strategy of GLAMS should be of great benefit in investigating differential changes in response to disease states from a glycoproteomics perspective, and has potential to contribute to biomarker discovery in many diseases.

## Supporting information

Figure S1-S9 and Scheme S1

Table S1. Identified sialoglycopeptide spectra for all human serum samples

Table S2. Distribution of site-specific sialoglycopeptides in human serum from colon cancer patients, liver cancer patients and normal people

## AUTHOR INFORMATION

### Author Contributions

H.Z., L. L., and P. G. W. conceived the project; H.Z. designed and performed the experiments, analyzed the data, and wrote the manuscript; S. W. synthesized the sialoglycopeptides; D. L. and L. D. prepared the serum glycopeptides for collision energy optimization; C. C., Y. L. and Z. W. synthesized the UDP-GalNAz; R. B. and K. L. collected and prepared the human serum samples; C. M., J. Y., L. L., and P. G. W. edited the manuscript. All the authors revised and approved the manuscript.

## ACKNOWLEDGEMENTS

We sincerely thank Dr. Wenfeng Zeng from the Key Lab of Intelligent Information Processing of Chinese Academy of Sciences for the help of parameter setting of pGlyco 3.0, Georgia Cancer Center Biorepository for providing human serum samples, and Georgia Research Alliance for financial support to purchase mass spectrometers. This work is supported by National Institute of Health (U01GM116263 and U54HL142019).

