## Supplementary material for "Identifying Sialylation Linkages at the Glycopeptide Level by Glycosyltransferase Labeling Assisted Mass Spectrometry (GLAMS)": Figure S1-S9 and Scheme S1

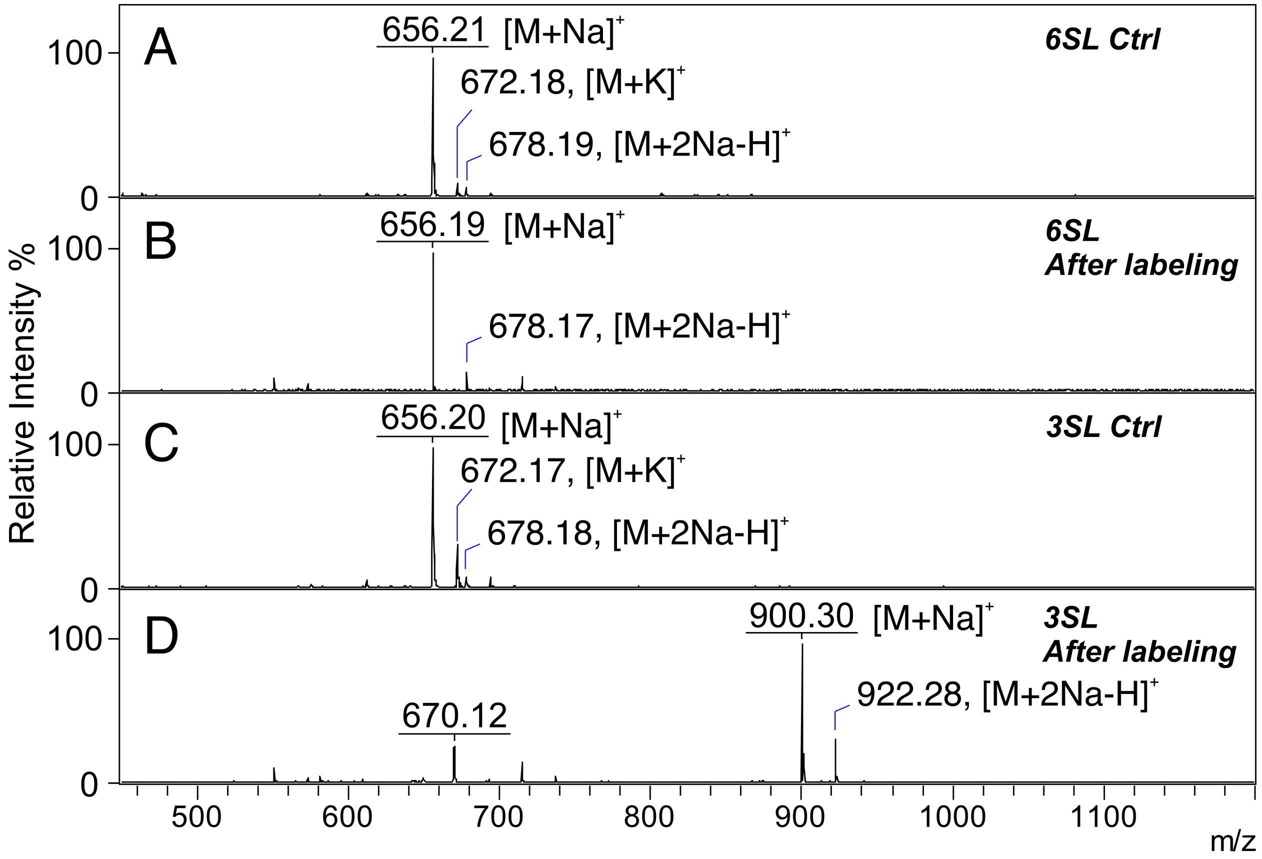

**Figure S1**. MALDI-MS spectra of 6SL (A) and 3SL (C) before enzymatic labeling and 6SL (B) and 3SL (D) after enzymatic labeling.

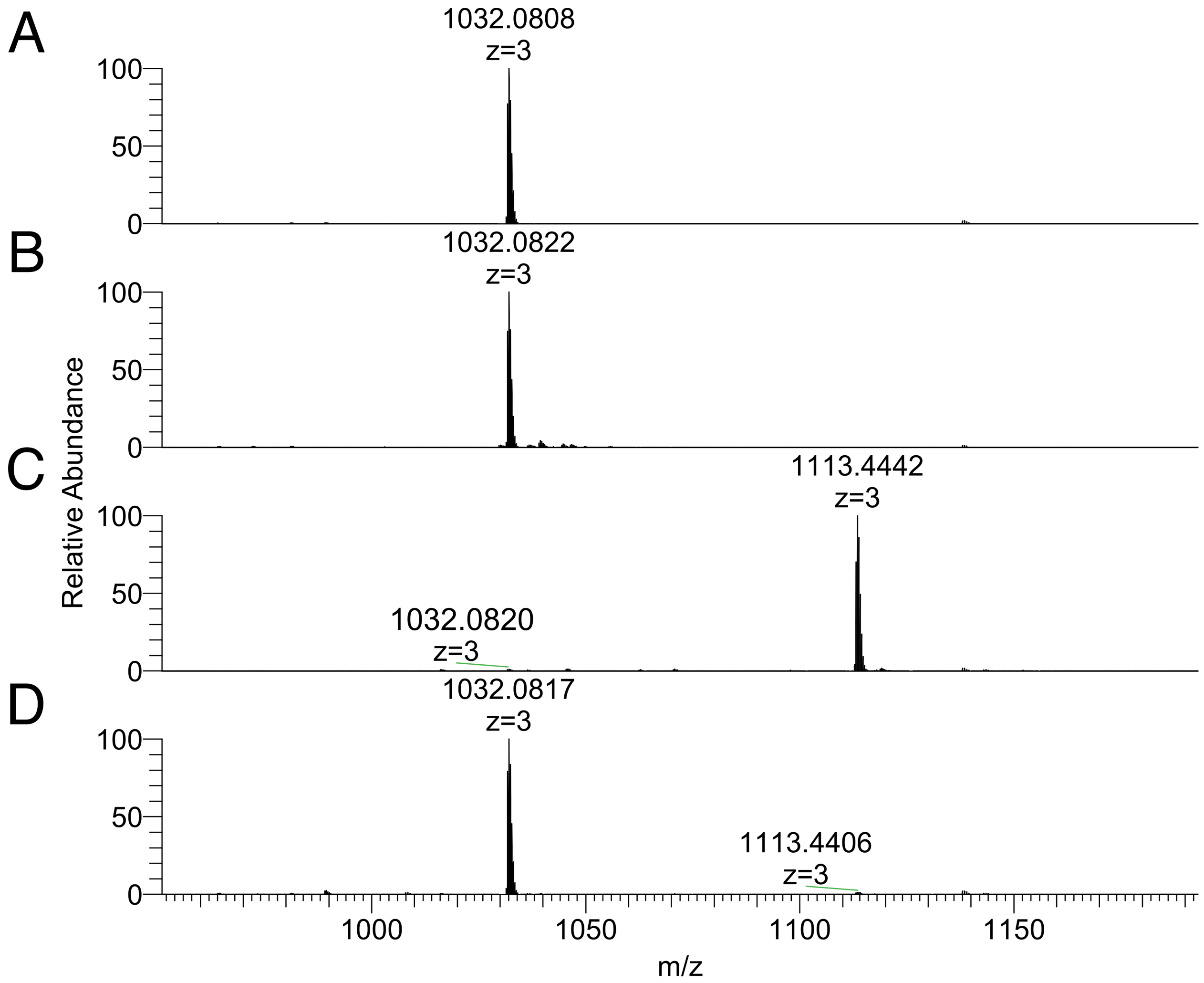

**Figure S2**. ESI-MS spectra of 23SP (A) and 26SP (B) before enzymatic labeling, and 23SP (C) and 26SP (D) after enzymatic labeling.

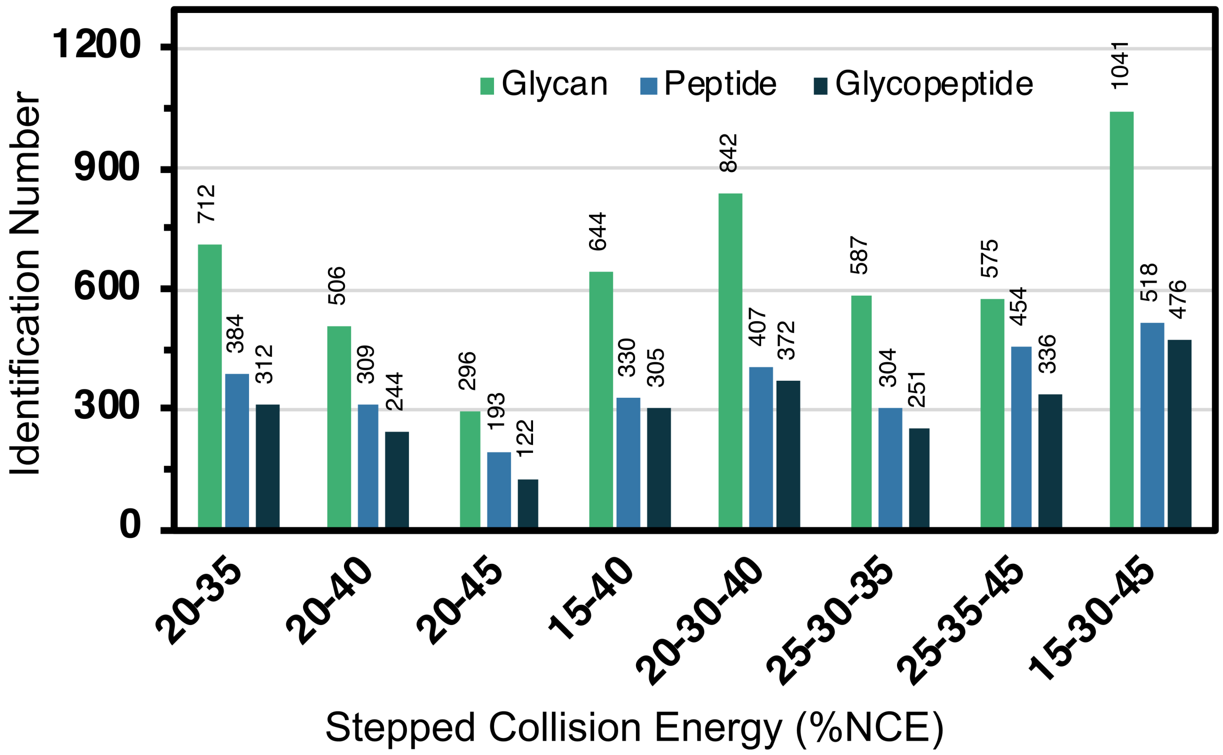

**Figure S3**. Identification number of N-glycopeptides in normal human serum under different collision energies. The identified glycans, peptides and glycopeptides were all filtrated with a FDR of 1%.

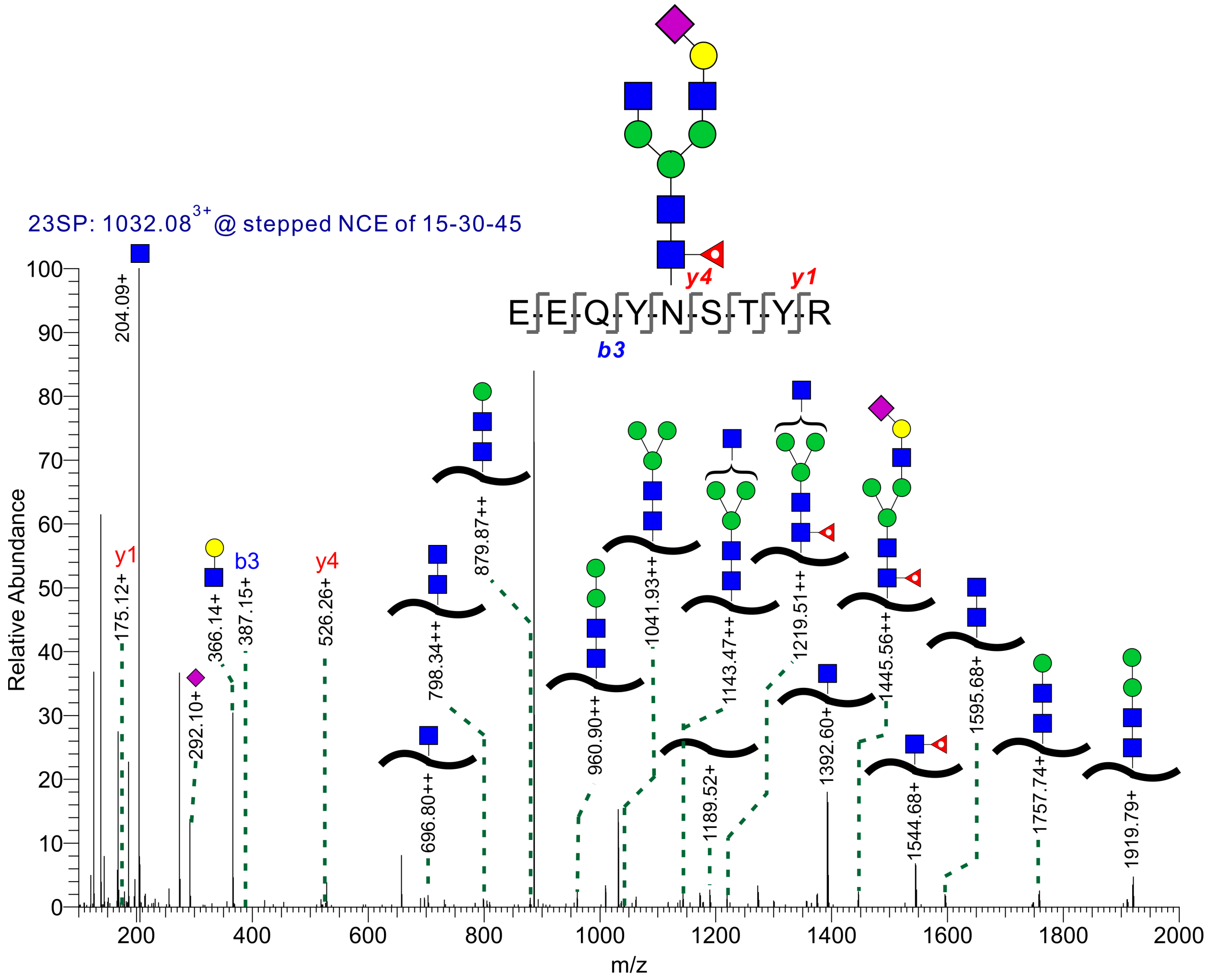

**Figure S4**. Tandem MS spectrum of 23SP with HCD fragmentation before enzymatic labeling.

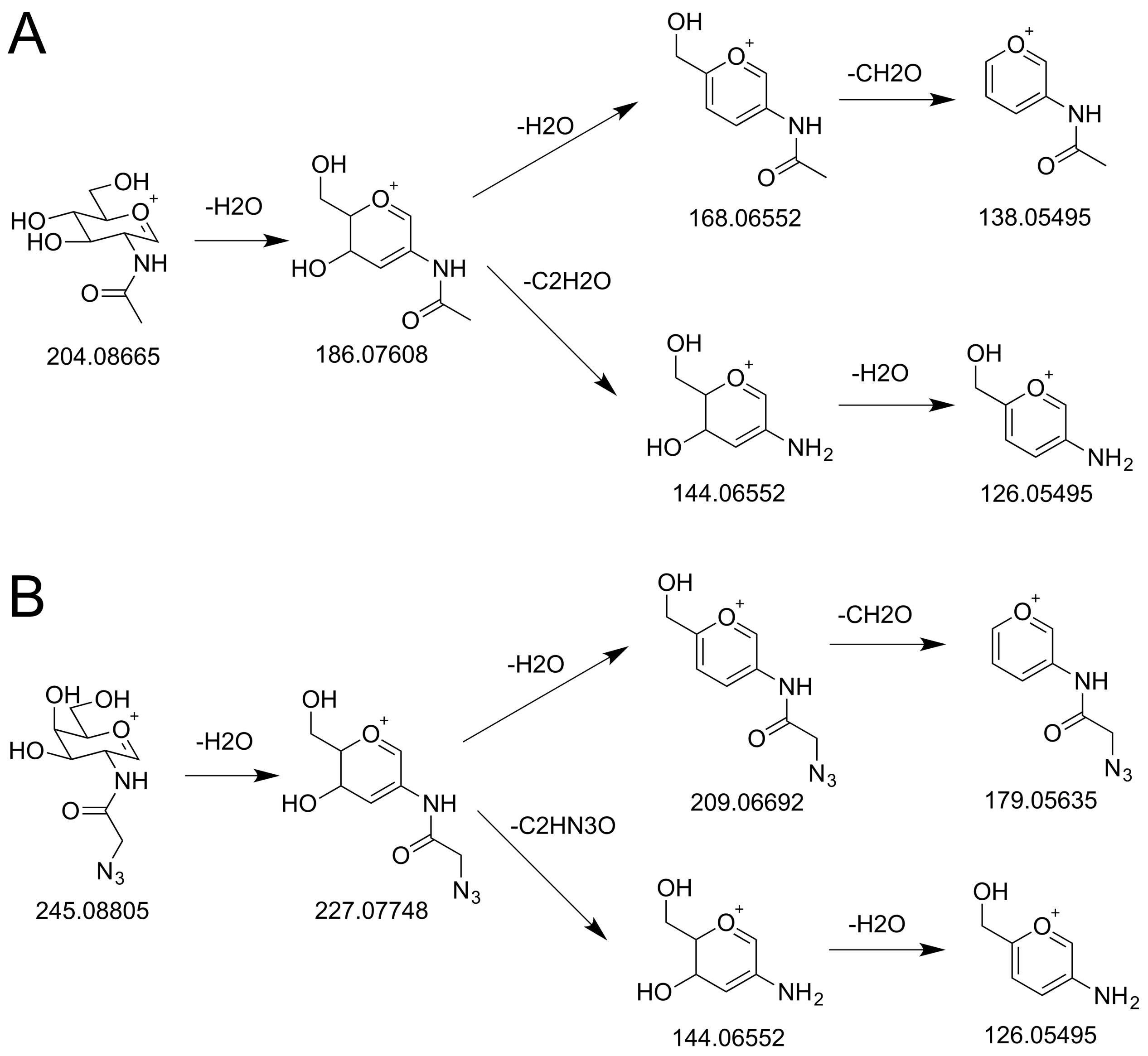

**Scheme S1**. Dissociation pathways for the GlcNAc (A) and GalNAz (B) in HCD mode.

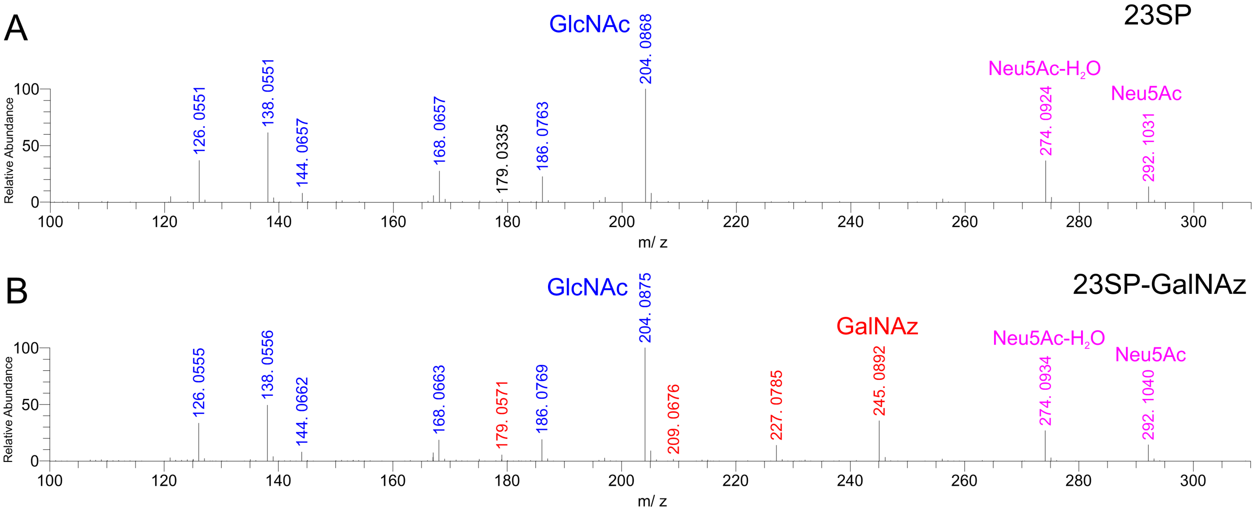

**Figure S5**. Zoomed-in spectra of 23SP before enzymatic labeling (A) and 23SP after enzymatic labeling (B).

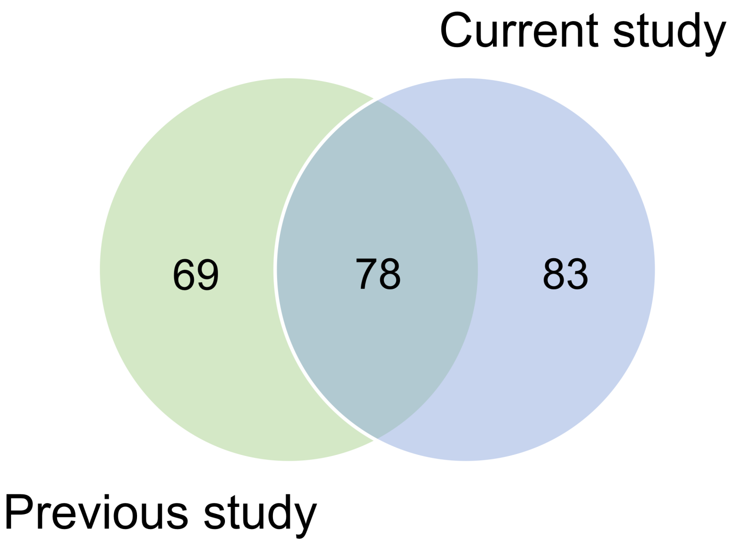

**Figure S6**. Venn diagram of identified glycoproteins from current study and previous study in which deglycosylation method was used.

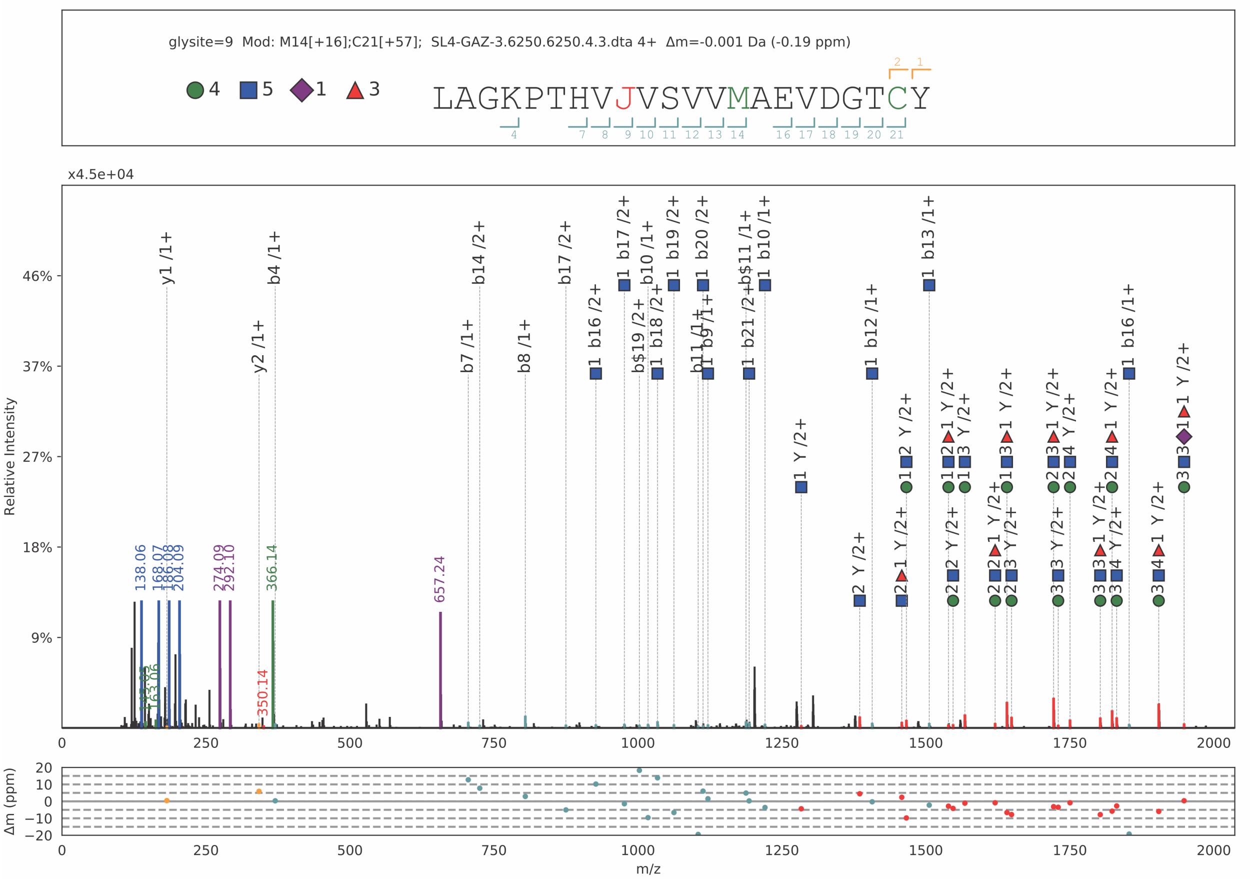

**Figure S7**. Representative annotated spectrum of α2,6-sialylated glycopeptide in the protein P01876 from liver cancer patients' serum. Green circles indicate Hexose, blue squares indicate GlcNAc, purple diamonds indicate Neu5Ac, and red triangles indicate Fuc.

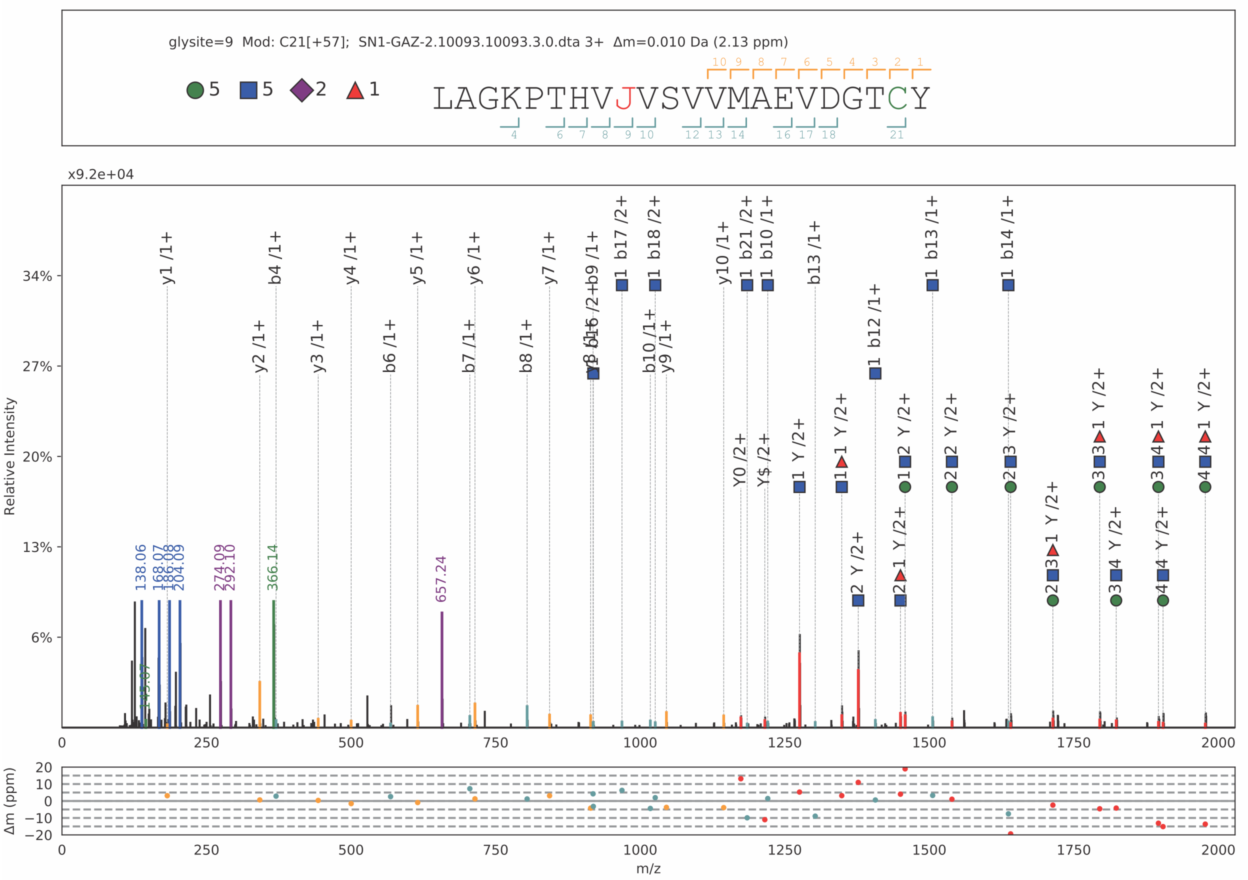

**Figure S8**. Representative annotated spectrum of α2,6-sialylated glycopeptide in the protein P01876 from normal human sera. Green circles indicate Hexose, blue squares indicate GlcNAc, purple diamonds indicate Neu5Ac, and red triangles indicate Fuc.

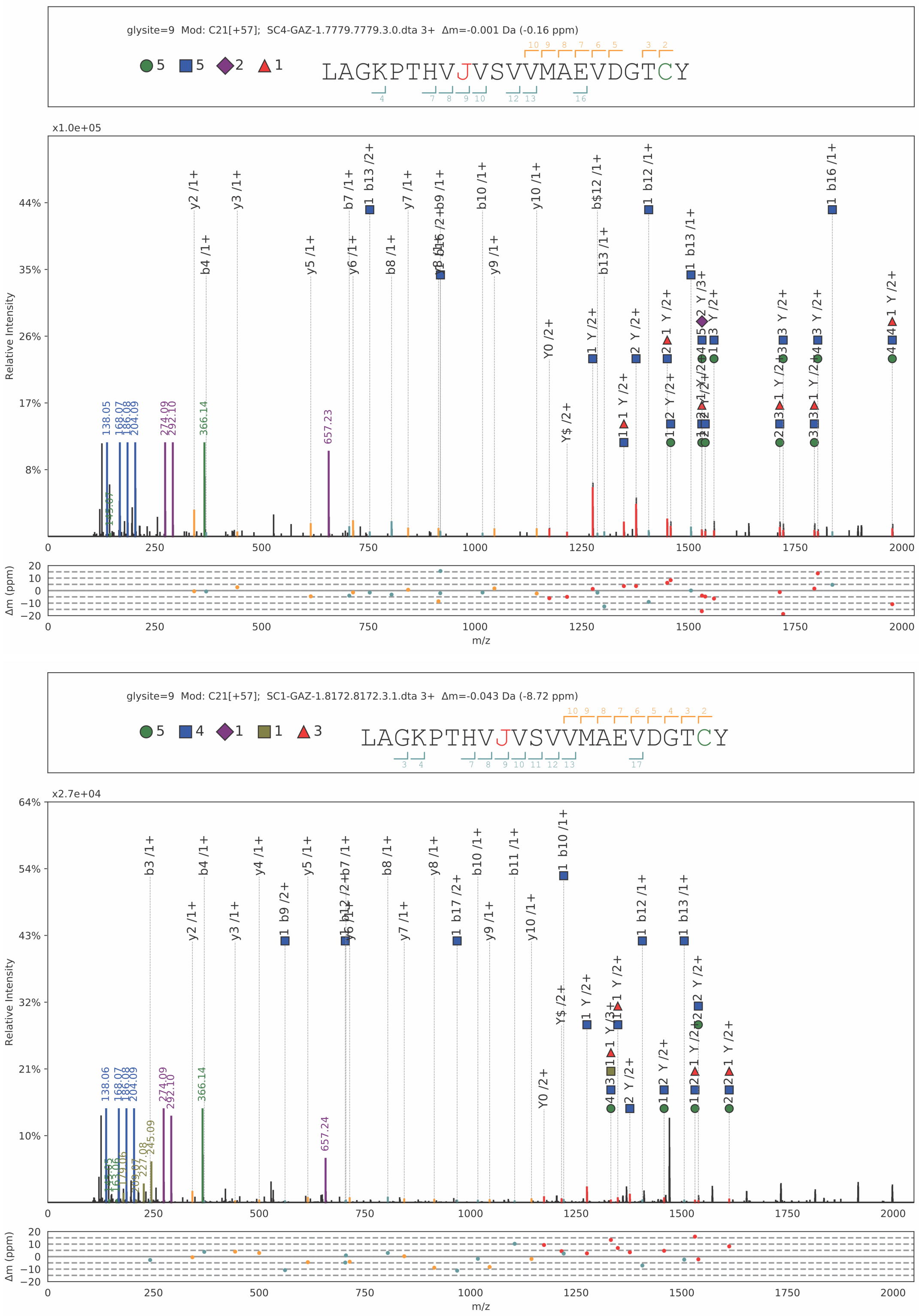

**Figure S9**. Representative annotated spectra of α2,6-sialylated glycopeptide (top) and α2,3-sialylated glycopeptide (bottom) in the protein P01876 from colon cancer patients' serum. Green circles indicate Hexose, blue squares indicate GlcNAc, olive squares represent GalNAz, purple diamonds indicate Neu5Ac, and red triangles indicate Fuc.
